# The interface of malignant and immunologic clonal dynamics in high-grade serous ovarian cancer

**DOI:** 10.1101/198101

**Authors:** Allen W. Zhang, Andrew McPherson, Katy Milne, David R. Kroeger, Phineas T. Hamilton, Alex Miranda, Tyler Funnell, Sonya Laan, Dawn R. Cochrane, Jamie L.P. Lim, Winnie Yang, Andrew Roth, Maia A. Smith, Camila de Souza, Julie Ho, Kane Tse, Thomas Zeng, Inna Shlafman, Michael R. Mayo, Richard Moore, Henrik Failmezger, Andreas Heindl, Yi Kan Wang, Ali Bashashati, Scott D. Brown, Daniel Lai, Adrian N. C. Wan, Cydney B. Nielsen, Alexandre Bouchard-Côté, Yinyin Yuan, Wyeth W. Wasserman, C. Blake Gilks, Anthony N. Karnezis, Samuel Aparicio, Jessica N. McAlpine, David G. Huntsman, Robert A. Holt, Brad H. Nelson, Sohrab P. Shah

## Abstract

High-grade serous ovarian cancer exhibits extensive intratumoral heterogeneity coupled with widespread intraperitoneal disease. Despite this, metastatic spread of tumor clones is non-random, implying the existence of local microenvironmental factors that shape tumor progression. We interrogated the molecular interface between tumor-infiltrating lymphocytes (TIL) and cancer cells in 143 samples from 21 patients using whole-genome sequencing, immunohistochemistry, histologic image analysis, gene expression profiling, and T- and B-cell receptor sequencing. We identify 3 immunologic response categories, which frequently co-exist within individual patients. Furthermore, epithelial CD8+ TIL were inversely associated with malignant cell diversity, evidenced by subclonal neoepitope elimination and spatial tracking between tumor and T-cell clones. Intersecting mutational signatures and immune analysis showed that foldback inversion genomic aberrations lead to worse outcomes even in the presence of cytotoxic TIL (n=433). Thus, regional variation in immune contexture mirrors the pattern of intraperitoneal malignant spread, provoking new perspectives for treatment of this challenging disease.

## Introduction

High-grade serous ovarian cancer (HGSC) exhibits the highest disease mortality among gynecologic cancers. Despite recent progress with synthetic lethal approaches exploiting homologous recombination deficiency through PARP inhibitors (Audeh et al., 2010; Gelmon et al., 2011), HGSC remains incurable with very poor survival rates. Characterized by profound genomic instability and extensive clonal diversity, HGSC often presents with disease widespread throughout the peritoneal cavity. Multi-site genomic studies have consequently revealed genomic measures of intratumoral heterogeneity as correlates to poor survival (Schwarz et al., 2015), and specific patterns of malignant cell spread within the peritoneal cavity (Bashashati et al., 2013). The physical distribution of clones across the peritoneal cavity is non-random, with the majority of sites exhibiting relative homogeneity and a minority of sites harboring phylogenetically diverse clones (McPherson et al., 2016). This raises the hypothesis that region-specific properties of invaded tissues, involving stromal and immunologic components of the tumor microenvironment, may modulate malignant cell invasion and expansion. In particular, observations of clonally diverse primary foci present in conjunction with distal clonally pure sites could indicate local immune privilege at sites with divergent clones and active immuno-selection at more clonally pure sites. Thus, to characterize the immuno-malignant interface in pre-treated HGSC, we carried out an unprecedented high resolution multi-modal and multi-site cohort-based study, assaying immune, stromal, and malignant cell compositions across peritoneal foci, revealing insights into malignant-immune interplay at the clonal level.

It is well established that HGSC tissues are subject to immune surveillance. Patients with abundant CD8+, CD4+, CD20+, and plasma cell tumor-infiltrating lymphocytes (TIL) are associated with favorable clinical outcomes (Zhang et al., 2003; Hwang et al., 2012; Nielsen et al., 2012; Kroeger et al., 2016). TIL can respond to and temporally track neoantigens (Wick et al., 2014), and through IFN-*γ* signalling, abolish fibroblast-mediated resistance to platinum compounds used in standard-of-care chemotherapy (Wang et al., 2016). Despite the prevalence of multi-site disease in HGSC, much of our understanding of the immune response in HGSC derives from single biopsies. Thus, the implications for disseminated disease are uncertain. Initial evidence from histologic images has revealed that lymphocyte abundance can vary between tumor foci in HGSC (Heindl et al., 2016). Furthermore, expression signatures of innate and adaptive immune responses are linked to patterns of metastasis (Auer et al., 2016). A single case report has described immunologic variation across relapse specimens (Jiménez-Sánchez et al., 2017); however, given the immunomodulatory effect of chemotherapy (Lo et al., 2017), it is unclear whether such variation exists before treatment. We contend that a cohort analysis linking spatial dynamics of malignant and immune cells via a pre-treatment multisite approach will provide context for interpreting such a remarkable and dynamic immunologic trajectory, along with emergent clinical trials investigating temporal immunologic response.

Recent discovery of prognostic mutational processes in HGSC through integrated analysis of point mutation, copy number and rearrangement features has indicated a prominent association between the prevalence of foldback inversions (FBI) and poor response to platinum-based chemotherapy (Wang et al., 2017). FBI-dominated tumors tend to be exclusive to homologous recombination deficient cases, and bear a distinct pattern of high level amplifications co-localized with foldback rearrangements typical of breakage-fusion-bridge processes (Campbell et al., 2010; Wang et al., 2017). FBI-dominated tumors comprise approximately 40% of all HGSC and identify the hardest-to-treat patients (Wang et al., 2017). As mutational processes are the likely mechanism for generating genomically diverse clones, understanding how the immune response modulates clones harbouring different mutational patterns may yield insight into the dynamics of disease progression.

We surmised that localized selective pressures imposed by the immune microenvironment shape the distribution of malignant clones during disease progression, impacting the substrate upon which chemotherapy acts. Thus, we systematically profiled the inter-relationship of clonal diversity, mutational processes and immunologic response across a cohort of patients and multi-region samples. Integrated genome sequencing-based clonal decomposition, transcriptome-based T-cell and B-cell receptor sequencing, multicolor immunohistochemistry, and histologic image analyses were applied. Our results integrate six orthogonal high resolution measurement assays, illuminating the landscape of cell-type interactions at the physical interface of malignant and immune cells across 143 tumor samples from 21 patients. We reveal an inverse relationship between immune infiltration and malignant clone diversity in HGSC, consistent with directional selection and immunoediting of genomically diverse clones. Furthermore, FBI mutational processes associate with poor survival even in highly infiltrated tumors, implying mutational processes as a potential mechanism for immune evasion. In aggregate, our findings illuminate the landscape of the immune-malignant interface and the evolutionary forces shaping tumor architecture prior to treatment in HGSC, providing context for analysis of temporal clinical trajectories.

## Results

### High-resolution multi-site profiling of immune and malignant populations in the HGSC tumor microenvironment

We assembled a cohort of 143 tumor samples from 21 HGSC patients. Multiple samples per patient were collected via pre-treatment primary debulking surgery from ovary, omentum, and other distant metastatic sites (except for relapse samples from patients 7, 11, and 23 (**Table 1**)). TIL densities were measured with multicolor immunohistochemistry (IHC); clonotype diversity in T- and B-cell populations with T- and B-cell receptor sequencing (TCR/BCR-seq); total mRNA gene expression from the 770-gene Nanostring PanCancer Immune Profiling Panel (Cesano, 2015) augmented with 39 molecular subtyping probes (Leong et al., 2015); mutational signatures and clonal diversity patterns of malignant cells from whole-genome (mean depth: 86X) and deep amplicon sequencing (mean depth: 16 278X, median number of loci: 188, **Table S1**); and 20X histologic images for local ‘ecosystem’ topologies and cell type colocalization (**Figure S1**). Clonal profiles and immune data (IHC, TCR/BCR-seq, or Nanostring) were obtained for 95 samples from 14 patients; data from all modalities were obtained for 46 samples from 14 patients. Details of sample acquisition and profiling are described in **Figure 1**A and **STAR Methods**.

**Table 1.** Studied patients and samples. Current disease status: NED, no evidence of disease; AWD, alive with disease; DOD, dead of disease. ^a^: BrnM 14 months, RPvM 33 months post-diagnosis; ^b^: Pv1, Rct1, Rct2 139 months postdiagnosis; ^c^: LOv1 14 months post-diagnosis

| Patient | Samples (index) | Anatomical samples (index) | Samples (archival) | Anatomical samples (archival) | Age at diagnosis (years) | Stage | Recurrence | RFS (months) | Status | OS (months) | BRCA status (clinical) |
| --- | --- | --- | --- | --- | --- | --- | --- | --- | --- | --- | --- |
| 1 | 6 | Om1,ROv1,R<br>Ov2,ROv3,R<br>Ov4,SBwl1 | 6 | ApC1,LFTB4,LO<br>vB2,RFTA16,R<br>OvA4,SBwlE4 | 71.7 | IIIC | No |  | NED | 71 | Screen negative |
| 2 | 4 | Om1,Om2,R<br>Ov1,ROv2 | 6 | LOvD3,OmA2,O<br>mB1,RFTC10,R<br>OvC2,ROvC4 | 75.5 | IIIC | Yes | 12 | DOD | 45 | Screen negative |
| 3 | 4 | Adnx1,Om1,<br>ROv1,ROv2 | 7 | CDSB1,LFTFim<br>C1,LOvC5,OmF<br>2,RFTA2,ROvA<br>7,SigClnDpE1 | 69.2 | IIIC | Yes | 25 | AWD | 73 | Screen negative |
| 4 | 5 | LPvSw1,ROv<br>1,ROv2,ROv<br>3,ROv4 | 4 | LOvSfcB2,LPvS<br>wC1,ROvA5,RP<br>vSwD1 | 52.9 | IIIA | Yes | 50 | AWD | 71 | Screen negative |
| 7<sup>a</sup> | 3 | BrnM,LOv1,<br>RPvM | 10 | BwlImA6,BrnM<br>A1,LOvA10,LOv<br>A4,ROvC4,ROv<br>C5,ROvC6,RUtD<br>1,RUtD2,RUtD3 | 47.2 | IIIC | Yes | 8 | DOD | 52 | Screen negative |
| 8 | 4 | EpSigCln1,O<br>m1,Om2,RO<br>v1 | 6 | LFTC1,LOvC3,O<br>mA3,OmB4,RO<br>vB3,SigEpD3 | 62.2 | IIIC | No |  | NED | 65 | BRCA1 mutation<br>and unclassified<br>BRCA2 variant |
| 9 | 5 | LOv1,LOv2,O<br>m1,Om2,RO<br>v1 | 0 |  | 52.5 | IIIB | Yes | 5 | DOD | 32 | Unknown |
| 10 | 4 | ROv1,ROv2,<br>ROv3,ROv4 | 4 | LFTB2,OmC1,R<br>OvA4,ROvA9 | 73.9 | IIIC | No |  | NED | 59 | Unknown |
| 11<sup>b</sup> | 5 | LOv1,LOv3,P<br>v1,Rct1,Rct2 | 0 |  | 53 | IIIB | Yes | 32 | AWD | 174 | BRCA2 mutation |
| 12 | 5 | LOv1,LOv2,O<br>m1,Om2,RO<br>v1 | 0 |  | 62 | IIIC | Yes | 15 | DOD | 44 | Screen negative |
| 13 | 5 | Om1,RGrn1,<br>ROv1,ROv2,<br>ROv3 | 0 |  | 80 | IV | No |  | NED | 40 | Screen negative |
| 14 | 5 | Bld1,CDS1,L<br>Ov1,Om1,RO<br>v1 | 0 |  | 58 | IIIC | Yes | 7 | DOD | 36 | Screen negative |
| 15 | 5 | CDS1,LOv1,R<br>Ov1,ROv2,SB<br>wl1 | 0 |  | 61 | IIIC | No |  | NED | 38 | BRCA1 VUS |
| 16 | 5 | LOv1,LOvSfc<br>1,Om1,Ptn1,<br>ROv1 | 0 |  | 72 | IIIC | Yes | 23 | AWD | 35 | Screen negative |
| 17 | 5 | Om1,Om2,O<br>m3,Ov1,Ov2 | 0 |  | 56 | IIIC | Yes | 19 | AWD | 32 | BRCA2 and MUTYH<br>mutation |
| 18 | 4 | ROv1,ROv2,<br>ROv3,ROv4 | 0 |  | 56 | IIIC | Yes | 19 | DOD | 34 | Unknown |
| 19 | 4 | ROv1,ROv2,<br>ROv3,ROv4 | 0 |  | 59 | IIIA | No |  | NED | 32 | Screen negative |
| 20 | 3 | LOv1,Om1,R<br>Ov1 | 0 |  | 64 | IIIA | No |  | NED | 10 | Unknown |
| 21 | 7 | LOv1,LOv2,R<br>Ov1,ROv2,R<br>Ov3,ROv4,R<br>Ov5 | 1 | LFT1 | 79 | IIIC | Yes | 4 | DOD | 45 | Screen negative |
| 22 | 8 | LFT1,LIN1,L<br>Ov1,LOv2,LO<br>v3,LOv4,LPa<br>N1,ROv1 | 0 |  | 73 | IIIC | Yes | 22 | AWD | 75 | Rare BRCA2 variant<br>(2680G>A), likely<br>benign |
| 23<sup>c</sup> | 3 | LOv1,Om1,O<br>m3 | 0 |  | 65 | IIIC | Yes | 9 | DOD | 75 | Screen negative |

**Figure 1.**
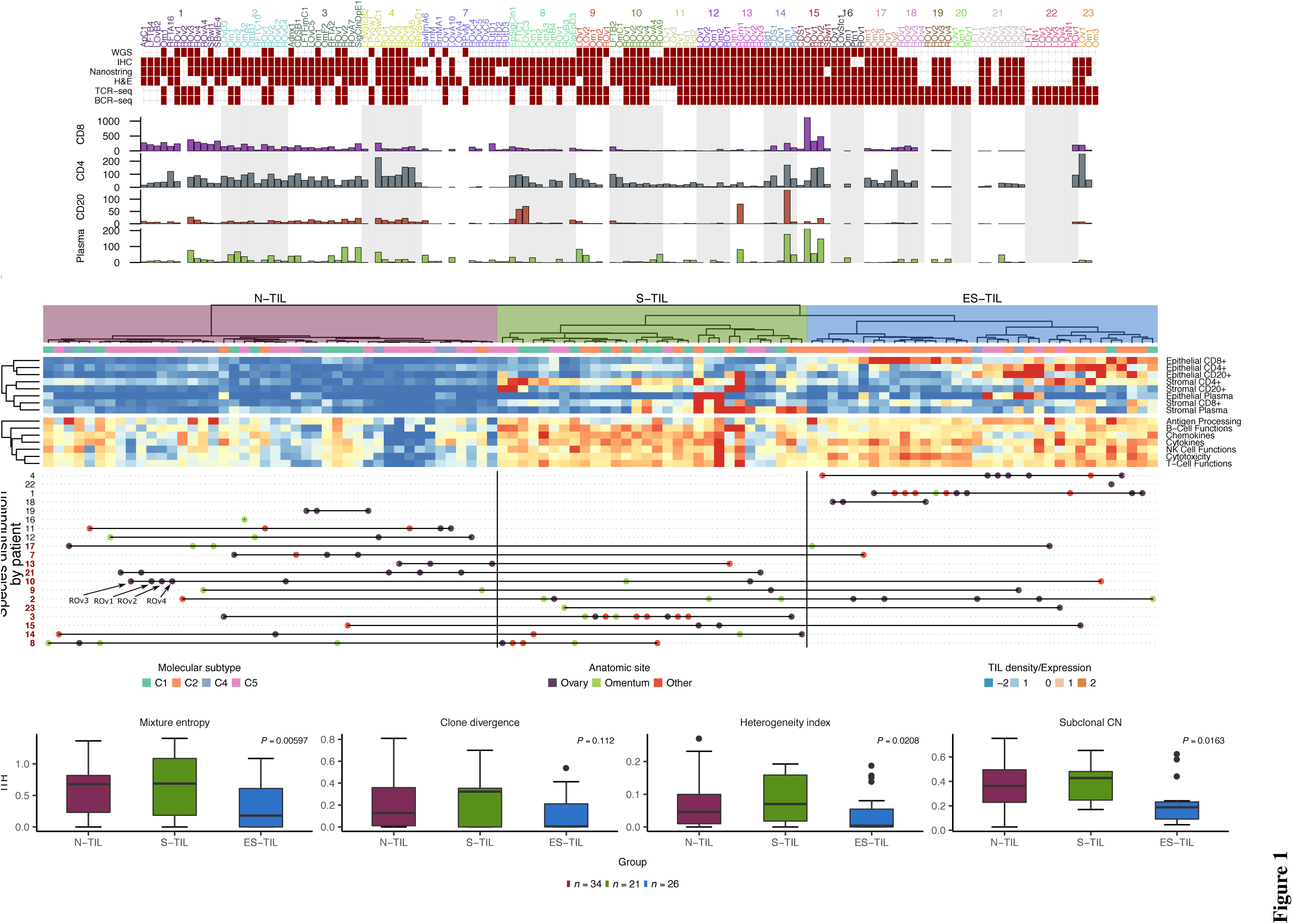
Sample TIL profiles and ITH. (A) Experimental matrix and IHC-based TIL densities. (B) Top: Epithelial and stromal TIL densities. Middle: Median expression for select immune pathways in Cesano (2015). Heatmap standardized and clipped between -2 and 2. Dendrogram computed from TIL densities with Ward’s method on L2-distance. Bottom: Sample labels. Patients in *≥* 2 clusters colored red. Only samples with both IHC and Nanostring data are shown. (C) ITH by TIL cluster. See also **Figure S1,2,4,5** and **Table S1-2**.

### Intrapatient variation in immune infiltration across peritoneal sites

We began by using IHC measurements to examine the degree of variation in immune composition across specific foci within patients. We profiled 119 tumor samples from 20 patients with multicolor IHC for CD8+ T cells (CD3+CD8+), CD4+ T cells (CD3+CD8-), CD20+ B cells (CD20+), and plasma cells (CD79a+CD138+) (with all but one patient surveyed at multiple sites). CD8+ T cells were the most abundant TIL type (1.16-1125.65 cells/HPF, median: 63.76), while CD20+ B cells were the rarest (0-136.77 cells/HPF, median: 2.74). Densities of all TIL types were correlated (**Figure S2**A), with extensive variation across the cohort (**Figure 1**A). However, no significant differences in TIL densities were observed between samples from ovarian, omental, or other peritoneal sites (**Figure S2**B) suggesting variation was independent of anatomic site.

Given the wide distribution of TIL densities across the cohort, we asked if samples could be grouped using TIL densities as features. Hierarchical clustering revealed 3 major TIL subtypes: N-TIL: tumors sparsely infiltrated by TIL; S-TIL: tumors dominated by stromal TIL; and ES-TIL: tumors containing both epithelial and stromal TIL (**Figure 1**B, **Table S2**). Gene expression values for several immune-associated pathways, including cytotoxicity, cytokines, and T- and B-cell associated genes, were comparable between S-TIL and ES-TIL, and lower in N-TIL, based on orthogonal Nanostring probe counts (**Figure 1**B). Furthermore, our immune groups mapped to previously described gene expression subtypes of HGSC: C1/mesenchymal, C2/immunoreactive, C4/differentiated, or C5/proliferative (**STAR Methods**) as defined by signatures from Leong et al. (2015). C4 and C5 tumors contributed to the bulk of N-TIL tumors (*p <* 1e-5, Fisher’s exact test), while C1 and C2 tumors were overrepresented in S-TIL (*p <* 0.01, Fisher’s exact test) and ES-TIL (*p <* 1e-5, Fisher’s exact test) tumors, respectively (**Figure 1**B), supporting the notion that previously reported HGSC gene expression classes are likely patterned by immune cell content. For 8/20 patients, only one class was observed (4/20 were N-TIL only; 4/20 were ES-TIL only; 0/20 were S-TIL only). The remaining 12/20 patients harbored tumors from more than one cluster (**Figure 1**B). N-TIL + S-TIL (5 patients) was a common combination, while N-TIL + ES-TIL was observed twice and S-TIL + ES-TIL only once. Four patients were represented in all 3 clusters. Thus, the distribution of samples across the 3 clusters indicates that immune infiltration profiles of tumor sites within a patient differ in approximately half of HGSC.

### Intratumoral heterogeneity is lowest in tumors with high epithelial lymphocyte infiltration

With evidence of immune response variation within intrapatient foci, we set out to determine the relationship between TIL subtypes and patterns of clonal diversity in malignant cells. Using whole genome sequencing from an index sample set of cryopreserved tissues (n=66, from 14 patients; 31 from 7 patients previously described (McPherson et al., 2016), we ascertained somatic single-nucleotide variants (SNVs), allele-specific copy number changes, and rearrangements (**Table S2**). We performed targeted deep amplicon sequencing on 97 tissue samples from these patients (including additional formalin fixed samples) to calculate clonal phylogenies and relative clonal composition of each sample (**STAR Methods**, **Figure S3**, **Figure 2**). We then related patterns of malignant clone composition with N-TIL, S-TIL and ES-TIL. Within each patient, tumors from the same TIL subtype did not have significantly higher clonal similarity than those from different subtypes (*p >* 0.3, permutation test, **Figure S4**B). For example, samples Om1 and Om2 from patient 17 were composed of identical tumor clones, and likewise Ov1 and Ov2 had comparable clonal composition (**Figure 2**); however, Om1 and Ov2 were ES-TIL whereas Om2 and Ov1 were N-TIL (**Table S2**). Thus, TIL subtype was likely not solely attributable to tumor clones.

**Figure 2.**
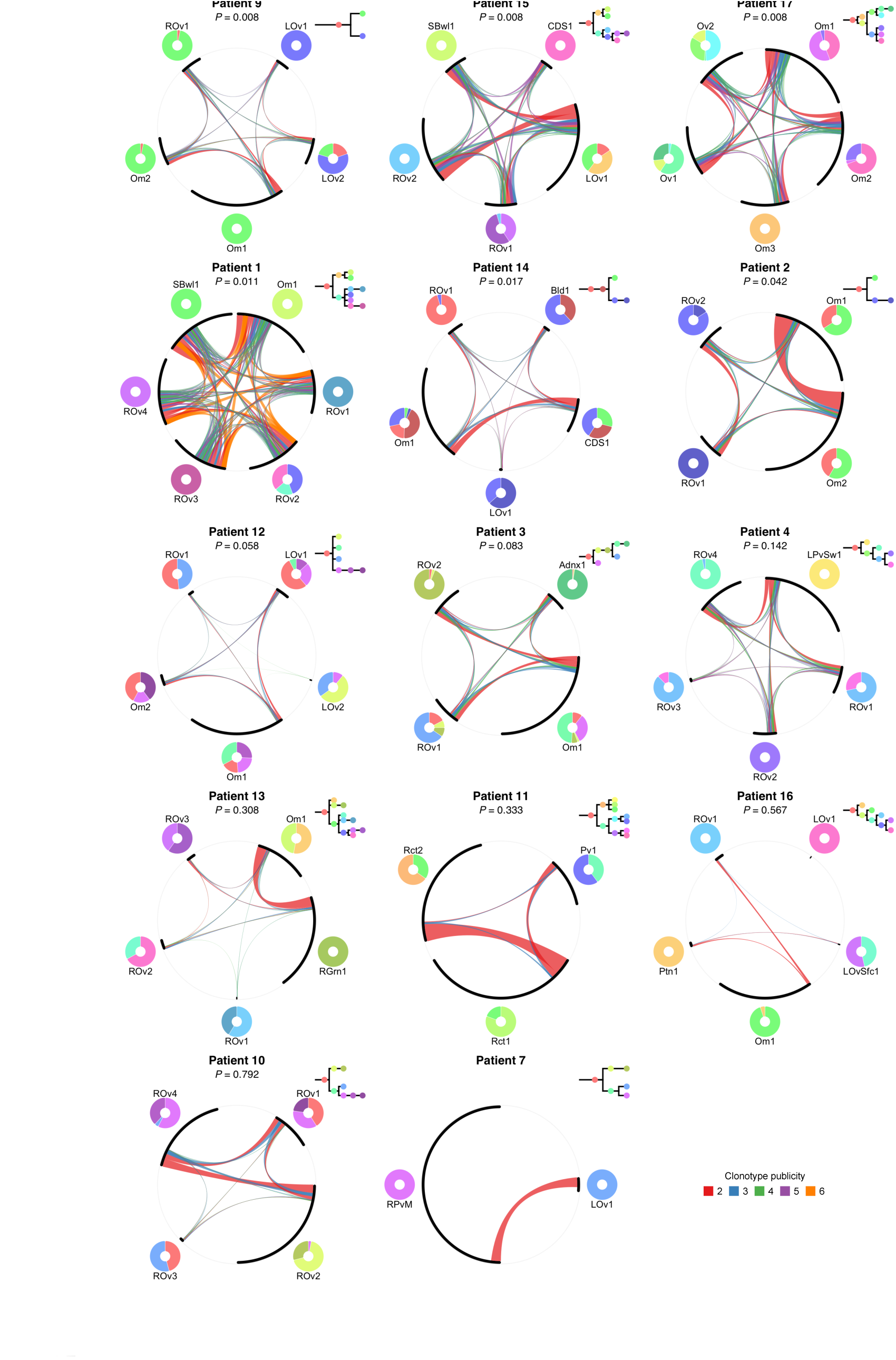
Malignant clone composition and TCR clonotype repertoires. Chords denote shared clonotypes, width proportional to number of clonotypes, colored by number of associated samples. Arc length proportional to number of clonotypes. Tumor clone composition and phylogenies shown next to chord diagrams. Uncorrected Mantel’s test *p*-values between TCR repertoire and clonal dissimilarities shown. See also **Figure S3**.

For each sample, we then summarized clonal composition into 3 continuous valued measures of complexity: mixture entropy - the mixture distribution of clones present within a sample; clone divergence - the maximum phylogenetic distance between clones present within a sample (McPherson et al., 2016); and heterogeneity index - the mean phylogenetic distance between a randomly selected pair of clones within a sample (weighted by abundance). In addition, we computed an orthogonal measure from WGS directly based on clonal inference from allele-specific copy number analysis (**STAR Methods**). We note that all four measures of ITH were correlated (all *p <* 0.05, significance of Spearman *ρ*), and none of the clonal measures were confounded by tumor purity (all *p >* 0.2, **Figure S4**A). We compared distributions of these measures as a function of the three immune classes described above. Strikingly, ES-TIL samples exhibited lower levels of all 4 measures than S-TIL and N-TIL (**Figure 1**C). Accordingly, clonally pure tumors had the highest epithelial CD8+ TIL densities (**Figure S4**C), establishing a negative association between epithelial TIL densities and the clonal complexity of malignant cells in HGSC.

### Epithelial CD8+ TIL are associated with subclonal neoepitope elimination

The negative association between epithelial TIL densities and malignant clone diversity implies at least two nonmutually exclusive scenarios: (1) clonally complex tumors may suppress the development of a TIL-rich microenvironment, or (2) in the presence of high epithelial TIL density, clones may be subject to immune-mediated purifying selection. In the latter scenario, subclonal (non-ancestral) neoepitopes might be subject to elimination. We computationally identified candidate neoepitopes from nonsynonymous somatic SNVs (4-123 nonsynonymous SNVs per sample, median: 40, **Table S3**, **STAR Methods**) in order to ascertain patterns of neoepitope elimination as a function of TIL. Putative neoepitopes were categorized as clonal or subclonal through phylogenetic analysis (**STAR Methods**). We quantified neoepitope elimination by comparing the observed neoepitope rate in each sample to the expected rate based on data from an independent cohort of 121 primary HGSC samples (**STAR Methods**, as per Rooney et al. (2015)). As ES-TIL tumors are primarily distinguished by high epithelial CD8+ densities (**Figure 1**B), we focused on the relationship between neoepitope elimination and epithelial CD8+ TIL. Within patients, samples with higher epithelial CD8+ density exhibited higher levels of subclonal neoepitope elimination (*p* = 0.01, generalized linear mixed model, **STAR Methods**) but not clonal neoepitope elimination (*p >* 0.1), suggesting that subclonal neoepitopes are immune-targeted in samples with high epithelial CD8+ TIL. This effect was restricted to epithelial CD8+ TIL, as no association was observed between clonal or subclonal neoepitope elimination and stromal CD8+ TIL density (all *p >* 0.3, generalized linear mixed model). Thus, samples with high epithelial CD8+ TIL show evidence of immune editing of subclonal neoepitopes, implying purifying selection away from malignant cell diversity.

### Regional variation in T-cell clonotypes tracks with the spatial distribution of tumor clones

We next asked whether T- and B-cell clonotypes associate with tumor clones. We applied TCR *β*-chain and BCR heavy chain sequencing to total RNA from 95 samples (21 patients, **Figure 1**A) to identify the clonotype-level composition of T- and B-cell populations in each sample. After quality control, 50 543 unique TCR (4-2714, median: 394 per sample) and 128 897 BCR clonotypes (6-6849, median: 832) were identified from 30 908 627 TCR (10 553-881 207, median: 343 076) and 45 356 059 BCR (28 098-1 166 270, median: 476 644) reads (**Table S2**, **STAR Methods**). TCR diversity was strongly correlated with IHC-based CD8+ and CD4+ TIL densities (all Spearman *p <* 1e-5, **Figure S2**C). Similarly, BCR diversity was significantly correlated with CD20+ and plasma cell densities (all Spearman *p <* 0.001, **Figure S2**D). C1 tumors had the most diverse TCR and BCR repertoires due to a higher proportion of rare clonotypes (**Figure S5**A,B). Only rarely were clonotypes observed in multiple patients (TCR: 0.04%, BCR: 0.03%).

We observed marked variation in intrapatient similarity for both TCR and BCR repertoires (**Figure 3**A-C, **Figure S2**E). Overall, intrapatient TCR and BCR repertoire similarities were correlated (Spearman *p <* 0.05), but with notable exceptions. Patient 15 had high TCR (ranked 3rd out of 21 patients) but not BCR similarity (15th), while patients 10 and 21 had high BCR (4th and 6th) but not TCR similarity (16th and 21st). The degree of intrapatient IHC- or Nanostring-based variation, though correlated to one another (**Figure S5**D), did not correlate with TCR or BCR repertoires (**Figure S5**C,E). For example, while 4 profiled samples from patient 10 (ROv1-4) had uniformly low expression of T-cell associated genes and CD8+ and CD4+ TIL densities (**Figure 1**B), they harbored distinct TCR repertoires (**Figure 3**C). Thus, higher resolution clonotype measurements carried additional information than could be ascertained by TIL densities alone.

**Figure 3.**
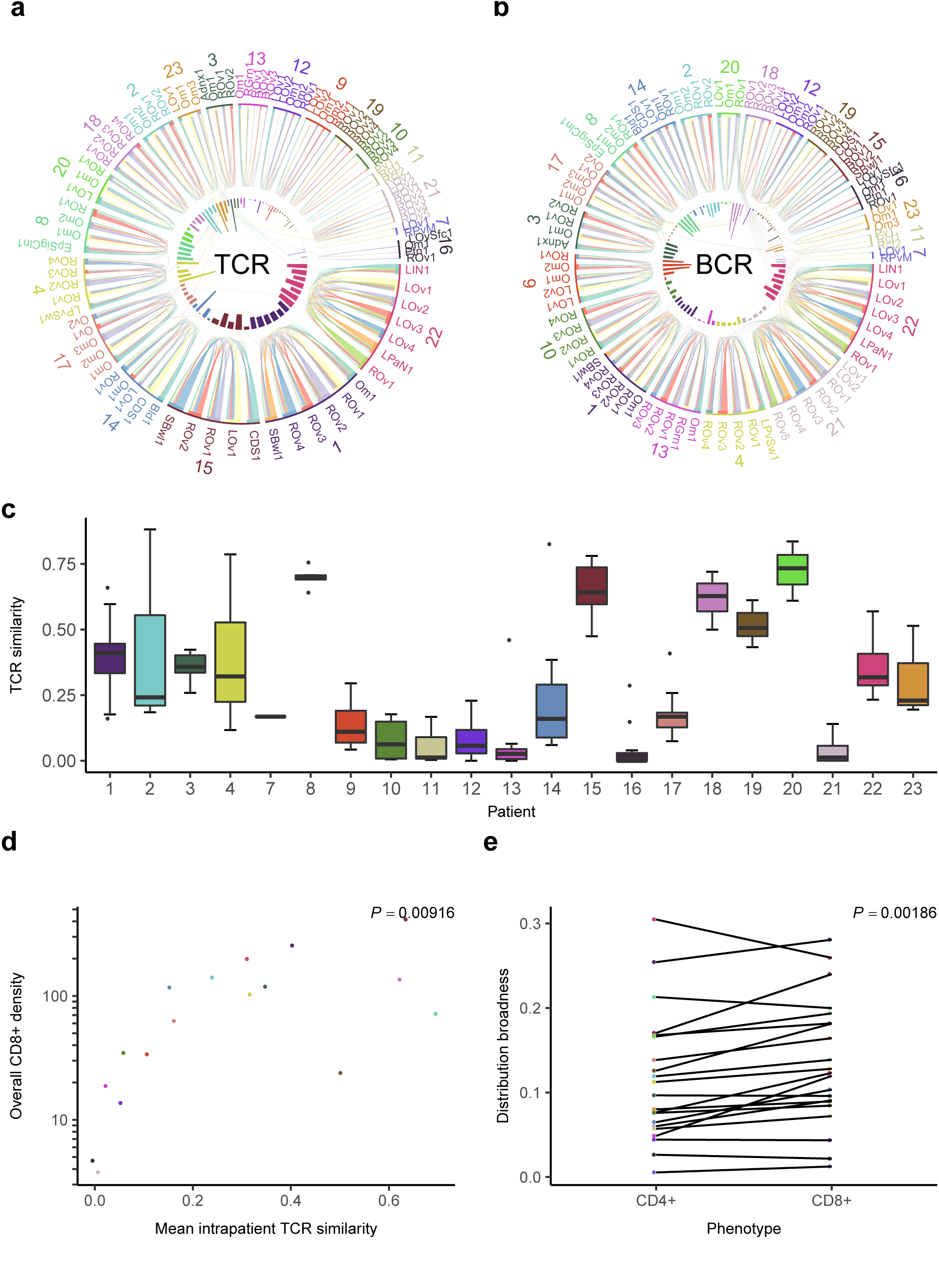
Spatial heterogeneity in TCR and BCR repertoires. (A+B) Shared clonotypes in TCR & BCR repertoires, shown as chords; width proportional to read count. Inner: number of clonotypes in each sample. (C) Pairwise TCR repertoire similarities. (D) Relationship between mean intrapatient TCR similarity and mean CD8+ TIL density. *P*-value of Spearman *ρ* shown. (E) Mean repertoire broadness for CD8- and CD4-type clonotypes in each sample. *P*-value from Wilcoxon signed-rank test. See also **Figure S2,5**, and **Table S2**.

Mean intrapatient TCR similarity was strongly associated with CD8+ (Spearman *p <* 0.01, **Figure 3**D) but not CD4+ TIL density (**Figure S2**F), suggesting that CD8+ TIL may be more broadly distributed (shared) across tumor sites compared to CD4+ TIL. To test this, we trained a classifier to separate TCRs as CD8- or CD4-type on the basis of V/J genes and physicochemical properties of the hypervariable domain (**STAR Methods**). The ratio of CD8/CD4-type TCRs was highly correlated with the ratio of CD8+/CD4+ densities by IHC (Spearman *p <* 0.001, **Figure S2**G). Corroborating our predictions, CD8-type TCRs were significantly more broadly distributed than CD4-type TCRs (*p <* 0.01, **Figure 3**E). In contrast, intrapatient BCR similarity was not significantly correlated with IHC-based CD20+ or plasma cell density (all Spearman *p >* 0.1).

Having established that TCR/BCR-based immune profiles vary across space, we asked how this variation is related to the spatial distribution of tumor clones. Pairwise T-cell repertoire similarity was significantly correlated with malignant clone composition similarity in 6 out of 13 patients (**Figure 2**). Importantly, this relationship was significant in 5/6 patients with the highest epithelial CD8+ TIL densities (patients 1, 2, 15, 17, and 9), implying that T-cell clonotypes spatially track with tumor clones in patients with high epithelial CD8+ TIL. This held when considering only major clonotypes (most abundant clonotypes constituting the top 50% of reads within each patient, significant in the same 6 patients), but not minor clonotypes (all other clonotypes, only significant in patients 2, 9, and 12), indicating that the most abundant clonotypes drove this effect. In contrast, pairwise BCR similarity was not significantly correlated with tumor clone similarity in any patient (**Figure S3**). None of the 4 ITH measures were significantly associated with TCR or BCR diversity (all Spearman *p >* 0.3, **STAR Methods**), indicating that diverse malignant populations do not recruit similarly diverse TIL repertoires.

### B-cell evolutionary dynamics recapitulate patterns of immune infiltration

Given the prognostic benefit associated with B cells in T cell-containing tumors (Nielsen et al., 2012; Kroeger et al., 2016), the absence of spatial tracking between B cells and tumor clones implies that B cells might recognize antigens with a more homogeneous spatial distribution. To first infer the extent of antigen-driven diversification and selection in B cells, we deciphered the evolutionary histories of B-cell lineages (clonotypes related through affinity maturation) using a Bayesian phylogeographic model (patients with temporal samples were excluded, **STAR Methods**). *Ab initio* comparison of framework (FR) and complementarity-determining regions (CDR) revealed that nucleotide substitution rates were higher in CDRs (median CDR/FR ratio: 2.67), consistent with the primary role of CDRs in antigen recognition (**STAR Methods**). Additionally, the mean CDR/FR ratio within each patient was positively correlated with epithelial CD8+ and CD4+ densities (all Spearman *p <* 0.05), suggesting that the presence of epithelial T cells promotes antigen-driven evolution in B cells. To explore this further, we asked whether selection operates to amplify specific clonotypes generated by antigen-driven diversification. If so, then phylogenetically related clonotypes should be more similarly expanded than distant clonotypes (**Figure 4**A). Using BCR transcript abundance as a measure of clonotype expansion, the degree to which this occurs in each lineage was quantified with *λ* (Pagel, 1999). Median *λ* for each patient positively trended with all epithelial TIL densities (all Spearman *p <* 0.054, **Figure 4**D), consistent with the notion that selective pressures act in a BCR-dependent manner to drive expansion of epithelial B cells. In contrast, median *λ* was not significantly correlated with stromal TIL densities (all Spearman *p >* 0.1).

**Figure 4.**
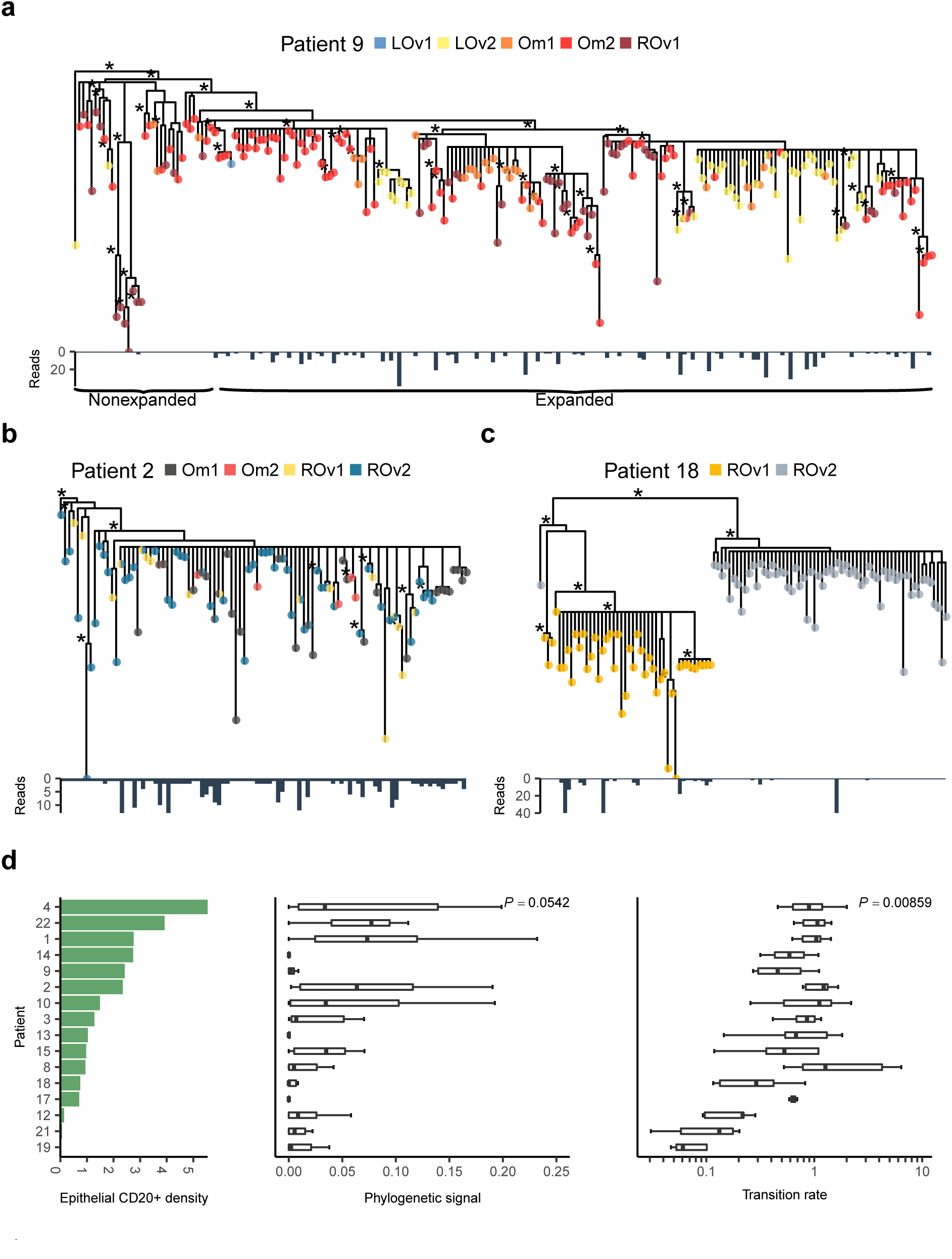
B-cell lineage phylogenies. (A+B+C) Lineages, showing (A) clade-specific expansion; (B) extensive inter-sample mixing; and (C) clade exclusivity between samples. Sample membership indicated at tree tips. Read counts indicated below each tree. Branch lengths proportional to substitution distance. Asterisks denote branching events with *>*80% probability. (D) Left: mean epithelial CD20+ density by patient. Middle: distribution of Pagel’s *λ*. Right: distribution of inter-sample transition rates. *P*-values of Spearman *ρ* between epithelial CD20+ density and median transition rate/*λ* shown.

Next, we evaluated the degree of spatial homogeneity in the B-cell response. To assess this, we examined sample-to-sample transition rates along each phylogeny (**STAR Methods**). In population genetics, sample-to-sample transitions are typically migration events, but in the context of rapidly evolving B cells they can also result from parallel somatic hypermutation occurring in different samples (von Büdingen et al., 2012). Phylogenies with transition values at both ends of the spectrum were observed. In some lineages, internal nodes gave rise to clonotypes found in multiple samples (**Figure 4**B), while other lineages contained sample-specific clades (**Figure 4**C). Median patient-level transition rates were positively correlated with epithelial and stromal CD20+ densities (all *p <* 0.01, **Figure 4**D) but not plasma cell densities (all Spearman *p >* 0.3).

Together, these findings imply that patients with high epithelial TIL densities mount spatially homogeneous, antigen-driven B-cell responses.

### Histologic interaction topologies of TIL and tumor cells

With multiple lines of evidence pointing towards coupled evolutionary dynamics between immune and malignant populations, we sought to verify that hotspots of TIL and malignant activity colocalize at the microscopic level. Exploiting the topological information from H&E slides interrogable with recently developed image processing algorithms (Yuan et al., 2012), we resolved patterns of spatial colocalization between TIL and cancer cells in 113 samples from 20 patients. First, we trained an algorithm to identify cancer cells, lymphocytes, and stromal cells within pathological tissue sections (**STAR Methods**). The relative fractions of cancer cells, lymphocytes, and stromal cells determined by the classifier were 0.27-0.86 (median 0.66), 0.01-0.38 (median 0.10), and 0.03-0.67 (median 0.23), respectively. Lymphocyte fraction was consistent with total TIL density measurements by IHC (Spearman *p <* 0.001).

Using cell locations, we applied spatial statistics (Getis and Ord, 1992) to elucidate hotspots of cancer cells and TIL (**Figure 5**A-C). We hypothesized that the extent of spatial colocalization between cancer and immune cells would be greatest in samples with high epithelial TIL density. To address this, we used 3 measures to quantify the degree of colocalization between lymphocyte and cancer cell hotspots (Nawaz et al., 2015): *f*_*C*_, the fraction of cancer cell hotspots that are lymphocyte hotspots; *f*_*I*_, the fraction of lymphocyte hotspots that are cancer cell hotspots; and *f*_*CI*_, fractional tissue area occupied by colocalized cancer-lymphocyte hotspots. All 3 measures were highest in ES-TIL samples (**Figure 5**D) and correlated with epithelial CD8+, CD4+, and CD20+ densities (all Spearman *p <* 0.01). Whereas cancer and TIL hotspots were mostly non-overlapping in N-TIL samples, they exhibited substantial overlap in ES-TIL, and, to a lesser extent, S-TIL samples (**Figure 5**A). Thus, our results provide evidence of localised interactions *in situ* between cancer cells and TIL in ES-TIL tumors.

**Figure 5.**
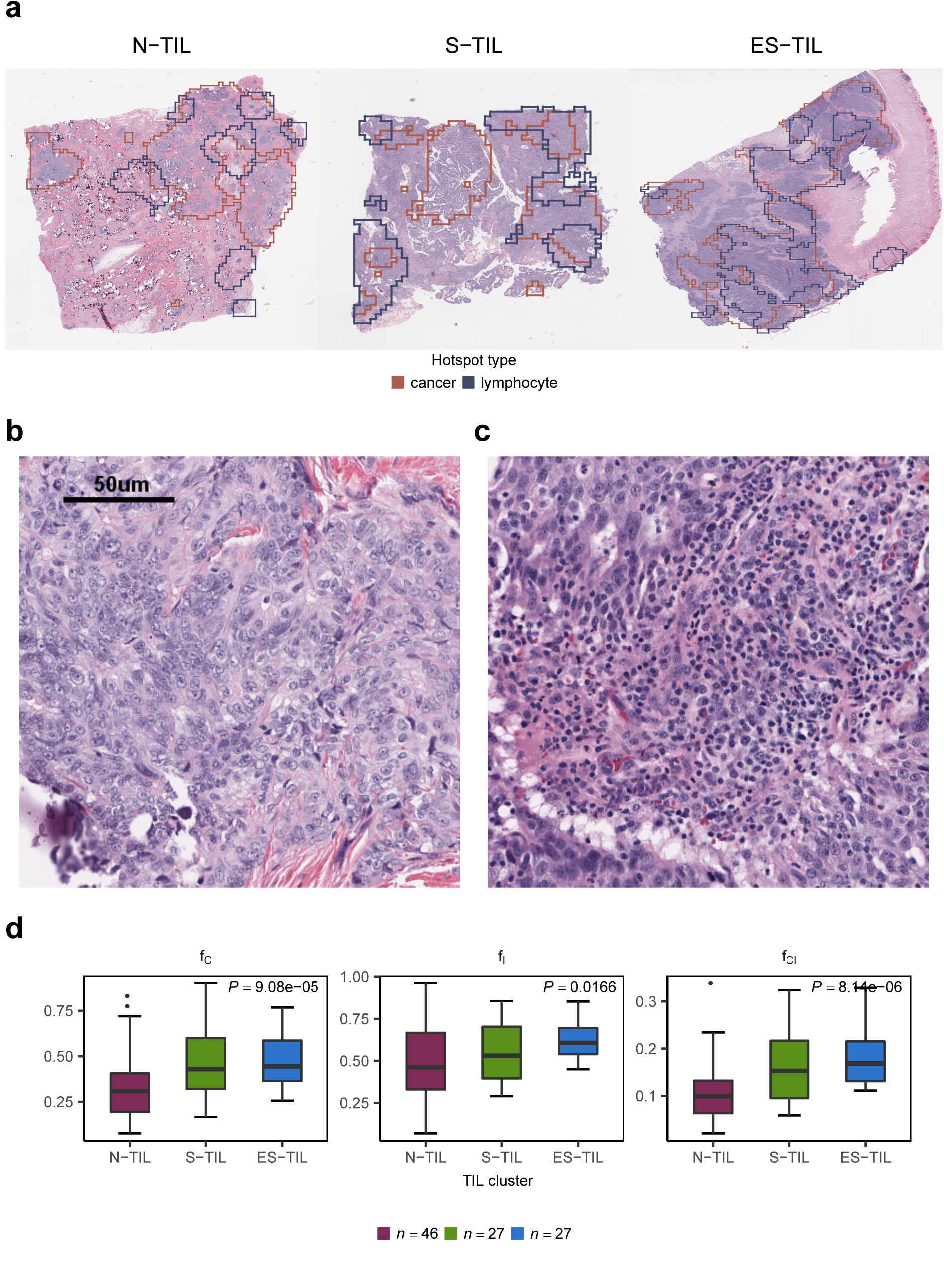
Cancer-lymphocyte hotspot colocalization. (A) Cancer cell and lymphocyte hotspots for representative N-, S-, and ES-TIL examples. (B) Histology of a cancer cell hotspot in a N-TIL sample. Tumor cells, but not TILs, are enriched. (C) Histology of a colocalized cancer and immune cell hotspot in an ES-TIL sample. Both tumor cells and TILs are enriched. (D) Comparison of cancer cell and lymphocyte hotspot colocalization between TIL clusters. *P*-values computed with Kruskal’s test. See also **Table S2**.

### Mutation signature correlates of immune activity

We previously profiled the mutational signatures present in HGSC, identifying 2 prognostically relevant subtypes: H-HRD (homologous recombination deficient) and H-FBI (foldback inversions) (Wang et al., 2017). Here, we investigated associations between those subtypes and observed immune properties. We applied a novel multimodal correlated topic model (MMCTM, Funnell et al., attached manuscript) to identify 6 SNV and 8 rearrangement signatures from 14 discovery cases and 194 additional single site ovarian cancer cases (132 from Wang et al. (2017), 62 from OV-AU in ICGC) (**Figure S6**A,B, **Table S4**). Hierarchical clustering by signature proportions identified 3 major clusters: one group (H-HRD) dominated by the point mutation signature associated with HRD (P-HRD), and two other groups (H-FBI-1 and H-FBI-2, collectively denoted H-FBI) characterized by a foldback inversion rearrangement signature (R-FB) associated with breakage-fusion-bridge cycles (Wang et al., 2017), but differing due to the age (P-AGE) and translocation signatures (R-TR) being more prominent in H-FBI-2 compared to H-FBI-1 (**Figure 6**A). No patients contained samples from multiple clusters (**Figure S6**C).

**Figure 6.**
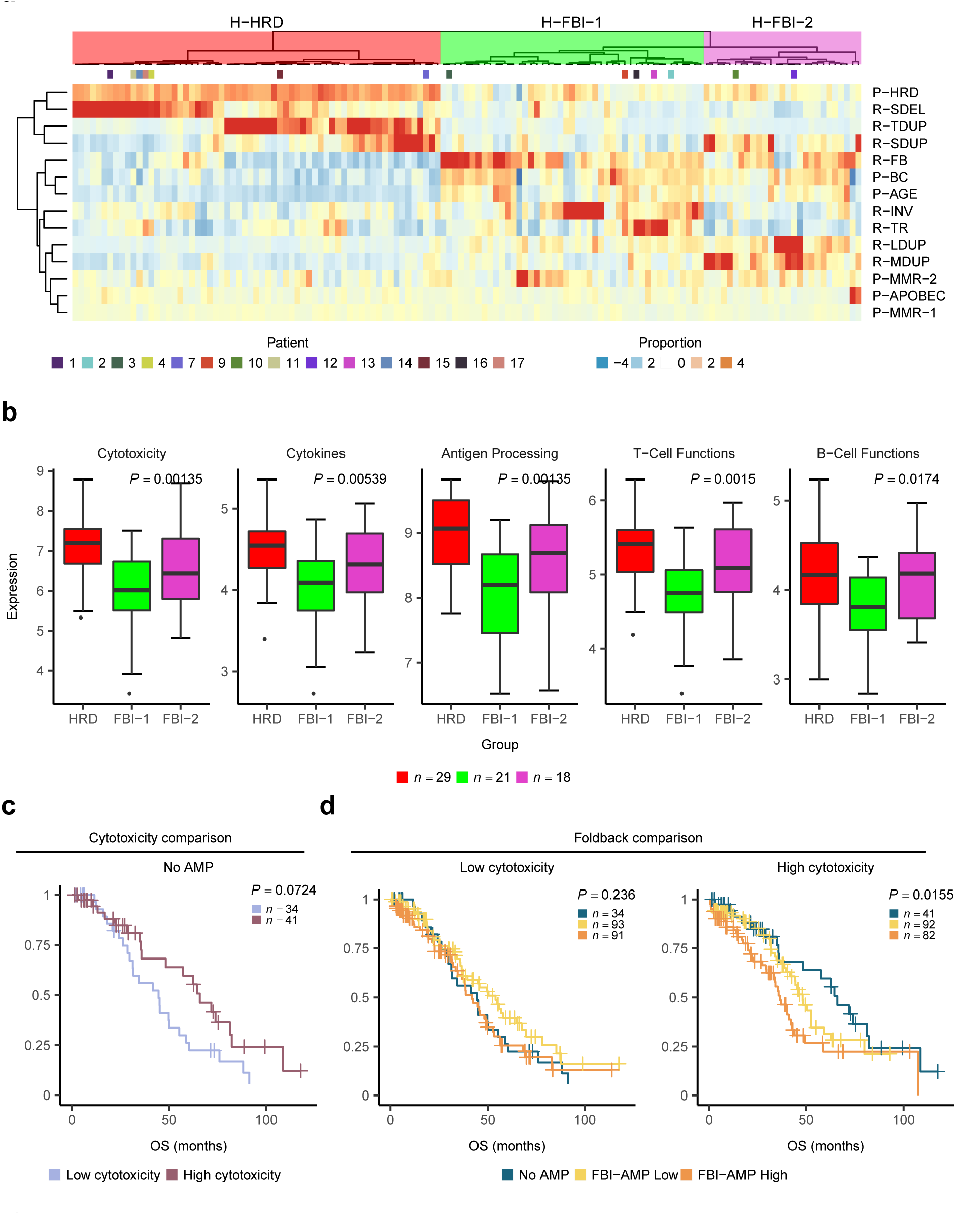
Mutation signatures and immunologic response. (A) Signature proportions for HGSC samples, standardized and clipped from -4 to 4. Dendrogram computed with Ward’s method on Pearson correlation dissimilarities. (B) Expression of select immune-associated pathways for mutational subtypes. Mean expression across samples used for discovery patients. *P*-values computed with Kruskal’s test and BH corrected. (C+D) Survival analysis of 433 TCGA samples. Differences in overall survival (C) between cytotoxicity subgroups for no HLAMP-tumors; (D) between FBI-HLAMP subgroups for tumors with low/high cytotoxicity separately. *P*-values computed with log-rank test. See also **Figure S6-7** and **Table S4-6**.

Using this grouping of samples, we asked how immune response characteristics co-segregated with mutational signatures. Overlaying Nanostring-based expression values of immune-associated pathways (Cesano, 2015) for 69 cases (**STAR Methods**) revealed that cytotoxicity, antigen processing, T- and B-cell markers were highest among H-HRD tumors (**Figure 6**B), concordant with similar findings in ER+ breast cancer (Smid et al., 2016) and among BRCA mutated tumors in HGSC (Nelson, 2015). Relative to H-HRD tumors, H-FBI-2 tumors had similar expression of immune markers, whereas H-FBI-1 tumors were depleted of these (**Figure 6**B). Corroborating these findings, whole-transcriptome differential expression analysis of OV-AU cases revealed that immune effector pathways were upregulated in H-FBI-2 relative to H-FBI-1, while no pathways were differentially expressed between H-HRD and H-FBI-2 (**Table S5**). Clustering by SNV or rearrangement signatures alone failed to recapitulate an association with immune signatures, emphasizing the importance of jointly considering both variant types.

Colocalized foldback inversions (FBI) and focal high-level amplifications (HLAMPs), thought to be reflective of breakage-fusion-bridge, have been associated with poor outcomes in HGSC (Wang et al., 2017). We asked whether immune activity could be used to further stratify FBI-enriched tumors into subgroups with distinct survival outcomes. Using array-based gene expression data for 433 ovarian cystadenocarcinoma cases from TCGA (Bell et al., 2011) (**Table S6**), we stratified cases into low and high cytotoxicity groups using the median value of the cytotoxicity signature defined above. In cases with colocalized FBI and HLAMPs, no significant effect of cytotoxicity on survival was observed (all *p >* 0.3, log-rank test, **Figure S7**). However, high cytotoxicity trended with increased overall survival in tumors with no HLAMP events (log-rank *p <* 0.1, **Figure 6**C). Doing the converse analysis, low FBI was associated with significantly longer overall survival among tumors with high cytotoxicity (log-rank *p <* 0.05, **Figure 6**D), but not low cytotoxicity (log-rank *p >* 0.2, **Figure 6**D). Hence, the prognostic benefit of low FBI or high immune activity is restricted to tumors with both properties.

## Discussion

Our results reveal profound intrapatient variation between immune microenvironments in HGSC, inversely linking spatial immune infiltrate properties to malignant cell diversity. Our data are consistent with active pruning of malignant cell diversity by TIL through subclonal neoepitope recognition. As such, we conclude that immune infiltrates act through purifying selection to shape patterns of malignant spread and clonal diversity in HGSC. The malignant-immune cell interaction comprises cell type-specific modes of clone tracking whereby B-cell clonotypes mount a spatially homogeneous response, while T-cell clonotypes track with malignant clones across peritoneal space. This pattern is enhanced in patients with the highest TIL densities, where high-resolution *in situ* histologic image analysis corroborated colocalization of malignant and immune cells. We note this does not exclude the possibility that T cells also recognize clonal neoepitopes (McGranahan et al., 2016; Jiménez-Sánchez et al., 2017). Our data indicate that co-registration of immune and malignant cell diversity may provide a new biomarker for patient or tissue sample stratification in clinical trials or retrospective cohort analyses.

We focused our attention on pre-treatment samples with the goal of understanding the interaction between malignant and immune cells in disease natural history up to diagnosis. As such, our study provides a landscape view of immune-malignant interaction, revealing the extent of variation and co-association prior to any treatment intervention. Our findings provide context for clinical trials investigating various classes of immunotherapy in ovarian cancer (e.g. immune-checkpoint blockade inhibition; adoptive T-cell transfer; and neoepitope vaccination). For instance, a recent case study tracking the immune response over time in a HGSC patient that followed a remarkable clinical trajectory (Jiménez-Sánchez et al., 2017) implies that spatial variation in immune response may play a major role in determining patient outcomes. Conceivably, the presence of even a single site experiencing relative immune privilege may be sufficient to engender resistant disease regardless of active immune responses mounted elsewhere. As a preliminary illustrative example, our TIL-subtype distribution across samples within patients could group patients into 3 categories: ES-pure, patients containing only ES-TIL samples; ES-mixed, patients containing both ES-TIL and N-TIL/S-TIL samples; and ES-none, patients containing no ES-TIL samples. Intriguingly, in this small cohort, ES-pure patients had better outcomes (median OS 71.3 months, 3/4 NED or AWD, 4/4 platinum sensitive) than ES-mixed and ES-none patients (median OS 41.6 and 44.9 months, 3/6 and 6/10 NED or AWD, 4/6 and 8/10 platinum sensitive, respectively).

Our data show for the first time a prognostic association between mutational signatures and immune response in HGSC. We show that a preponderance of foldback inversions, even in highly cytotoxic tumor microenvironments, are associated with the poorest outcomes. This suggests that, in contrast to mismatch repair deficiency in colorectal cancers (Le et al., 2015; Xiao and Freeman, 2015), FBI may generate genomic aberrations that impede immune surveillance even in the presence of cytotoxic TIL. Conversely, high immune response patients with low prevalence of FBI showed the most favorable outcomes.

The negative correlation between TIL and ITH underscores the clinical challenge of clonally heterogeneous tumors harboring the weakest TIL responses. These tumors may act as reservoirs of clonal diversity, providing immunologically sheltered havens from which treatment-resistant clones can emerge. While clonally diverse samples are the minority overall, most patients are not ES-pure (16/20) and unfortunately harbor at least one diverse tumor (McPherson et al., 2016). Given that clinical efficacy of PD-1 axis blockade hinges on pre-existing adaptive immunity (Melero et al., 2014; Herbst et al., 2014), these tumors may underlie the limited success of immunotherapy in HGSC to date (Homicsko and Coukos, 2015; Gaillard et al., 2016). Further studies will be necessary to evaluate how TIL-poor sites in ES-mixed patients respond to immunotherapy. If this obstacle can be surmounted, our findings hint at the tantalizing potential that immunomodulation of the tumor microenvironment may be able to hamper clonal diversification, reducing the likelihood of resistance to subsequent treatment.

As the cancer evolution field progresses towards a more rigorous understanding of the fitness of heterogeneous clones within disease spectra and over temporal dimensions (Lipinski et al., 2016), it is clear the external selective pressures imposed by the immune system must be considered as highly relevant factors. Here we show that high-resolution measurement of the immune microenvironment together with high-resolution clonal decomposition analysis is tractable and yields novel insight into forces shaping malignant cell diversity and intraperitoneal spread. Broadly disseminated intraperitoneal disease at diagnosis in HGSC remains a formidable clinical problem. Our study informs on how regional variation in immune response can control dissemination of clones and identifies critical tumor microenvironmental properties to exploit in future design of immuno-oncologic therapeutic strategies for HGSC.

## Author contributions

Study design, A.W.Z., R.A.H., B.H.N., S.P.S.; Writing, A.W.Z., B.H.N., S.P.S.; Manuscript review, A.W.Z., B.H.N., S.P.S., R.A.H., S.A., A.B-C., Y.Y., Y.K.W., A.B., W.W.W., C.B.G., C.d.S.; Data interpretation, A.W.Z., P.T.H., D.R.K., A.Mi., T.F., A.Mc., S.D.B., J.N.M., R.A.H., B.H.N., W.W.W., S.P.S.; Curation, D.R.C., D.L., A.N.C.W.; Formal analysis, A.W.Z., A.Mc., A.R., T.F., Y.K.W., A.B.; DNA/RNA extraction, J.L.P.L., W.Y.; IHC, K.M., D.R.K., S.L., A.W.Z.; TCR/BCR-seq, K.T., T.Z., I.S., M.R.M., R.M., H.F., A.H., Y.Y.; Nanostring, J.H.; Pathologic evaluation, A.N.K., C.B.G.; Supervision, S.P.S., B.H.N., R.A.H., D.G.H., W.W.W.; Visualization, A.W.Z., M.A.S., C.B.N.

## Acknowledgments

We thank Dr. Erick Matsen, Dr. Duncan Ralph, Kristian Davidsen, and Dr. Hossein Farahani for their advice regarding phylogeography. Funding was provided by the BC Cancer Foundation (S.P.S., B.H.N., R.A.H.), Canadian Cancer Society Research Institute (S.P.S., B.H.N., R.A.H.), Canada’s Networks and Centres of Excellence (B.H.N., R.A.H.), Canadian Institutes of Health Research (S.P.S., B.H.N., P.T.H.), Terry Fox Research Institute (B.H.N.), Cancer Research Society (B.H.N.), Genome Canada and Genome British Columbia (S.P.S., R.A.H.), Canada Research Chairs Program (S.P.S., D.G.H.), Michael Smith Foundation for Health Research (S.P.S., A.Mc.), and Vanier Canada Graduate Scholarship (A.W.Z.).

